# Examining Cognitive Performance in Mice using the Open-Source Operant Feeding Device FED3

**DOI:** 10.1101/2024.04.04.588157

**Authors:** Laura B. Murdaugh, Brieann Brown, Chin-Hui Chen, Cristina Miliano, Yuyang Dong, Starlina Shepard, Jason W. Putnam, Christine L. Faunce, Luis A. Natividad, Sujith Vijayan, Ann M. Gregus, Matthew W. Buczynski

## Abstract

Cognitive impairments are prevalent in various neurological disorders, including chronic pain conditions, and pose significant therapeutic challenges. Preclinical rodent models serve as valuable tools for investigating the underlying mechanisms of and treatments for cognitive dysfunction. However, factors such as stress, age, sex, and disease duration present challenges to reliably capturing cognitive deficits in rodents. Here, we present a comprehensive and high-throughput protocol utilizing the open-source operant Feeding Experimentation Device 3 (FED3) for assessing cognitive performance in mice. We developed a data pipeline to streamline data compilation and analysis, and established operating conditions for a six-test cognitive battery which can be completed in as few as 20 days. We validated our testing procedures using bilateral orbitofrontal cortical lesions to capture deficits in executive function, and demonstrated the feasibility of assessing cognitive function in aged mice of both sexes to identify genotypic and sex-specific effects. Overall, our findings demonstrate that the FED3 is a versatile tool for evaluating cognitive function in mice, offering a low-cost, high-throughput approach for preclinical studies of neurological disorders. We anticipate that this protocol will facilitate broader implementation of cognitive testing in rodent models and contribute to the understanding and treatment of cognitive dysfunction in neurological diseases.

## INTRODUCTION

Cognitive behaviors encompass a wide range of mental processes regarding how an organism integrates emotions, memories, and sensory information to make decisions on how to interact with others or their environment. Impaired or altered cognitive behaviors are a primary feature of numerous neurological disorders including addiction, anxiety, depression, Alzheimer’s, and pain^1–4^. Specifically, patients with chronic neuropathic pain^5,6^, including chemotherapy induced peripheral neuropathy^7^, report cognitive changes that are often under-treated by the traditional analgesics prescribed for these conditions^8,9^. For this reason, cognitive behavioral therapy^10,11^ has been widely studied as a non-pharmacological treatment approach and future therapeutic strategies are needed to specifically target cognitive dysfunction.

Preclinical rodent models provide an important tool for advancing our understanding of the mechanisms underlying cognitive dysfunction during chronic pain states and other neurological disorders. Previous work shows these conditions can result in disease-induced negative affect^12^ and deficits in motivated behavior, reflecting comorbidities observed in clinical populations^13,14^. However, using preclinical models to reliably capture clinical symptoms faces numerous experimental challenges. Cognitive performance in rodents is influenced by various non-cognitive factors including stress levels, age, handling history, and sex, which can negatively impact rodent cognitive performance and test reliability^15–17^. Furthermore, many inducible disease models transiently exhibit symptomatic expression (often days or weeks in duration), highlighting the need for cognitive testing paradigms that can be performed quickly.

Historically, spatial learning and memory tasks^18^ (e.g. Morris Water Maze^19,20^, Barnes Maze^16^) and recognition tasks (e.g. novel object recognition^21–23)^ have been implemented to measure cognitive function in large cohorts of mice^12,24–28^, including chemically-induced rodent pain models such as Chemotherapy Induced Peripheral Neuropathy^29–31^ and Complete Freund’s Adjuvant^32^ which exhibit a limited period of pain sensitivity. However, these paradigms rely on hippocampal-based spatial memory and the HPA-driven stress-influenced motivated behavior that provide an experimental challenge for evaluating executive function and other subdomains of cognitive behavior. Alternatively, operant tests^33–36^ using food as a reinforcer have been developed and implemented to evaluate such subdomains of cognitive and executive function including reinforcement learning and goal-directed behavior^37–43^. However, this approach often requires substantial animal training^44–47^ that may extend beyond the period of symptomatic expression in inducible disease models, imposing practical limitations to their use. Thus, preclinical evaluation of executive function and other cognitive impairments during chronic pain states requires approaches that are well suited for screening large number of mice in a short period of time^48^.

Here, we present a protocol for evaluating cognitive performance in mice using the Feeding Experimentation Device version 3^49–51^ that addresses many of these challenges (**Figure 1**). The FED3 is an open-source operant device that is suitable for use in a traditional mouse home-cage environment, reducing the influence of environment-induced stress on cognitive performance, while increasing the number of cognitive behavioral tests researchers can perform each week. We established operating conditions using the FED3 to perform an in-cage 6-test cognitive battery that includes tests capturing affective behaviors and executive function, and can be completed in as few as 20 days. Using bilateral lesions targeting the lateral and ventral orbitofrontal cortex (OFC), we validated that our testing procedures identified deficits in multiple aspects of executive function. Furthermore, we demonstrated that our test battery functions equally well in aged mice of both sexes. Finally, we developed an open-source workflow that streamlines data compilation and analysis to facilitate more comprehensive preclinical evaluation of cognitive dysfunction during chronic pain and other neurological disorders.

**Figure 1.**
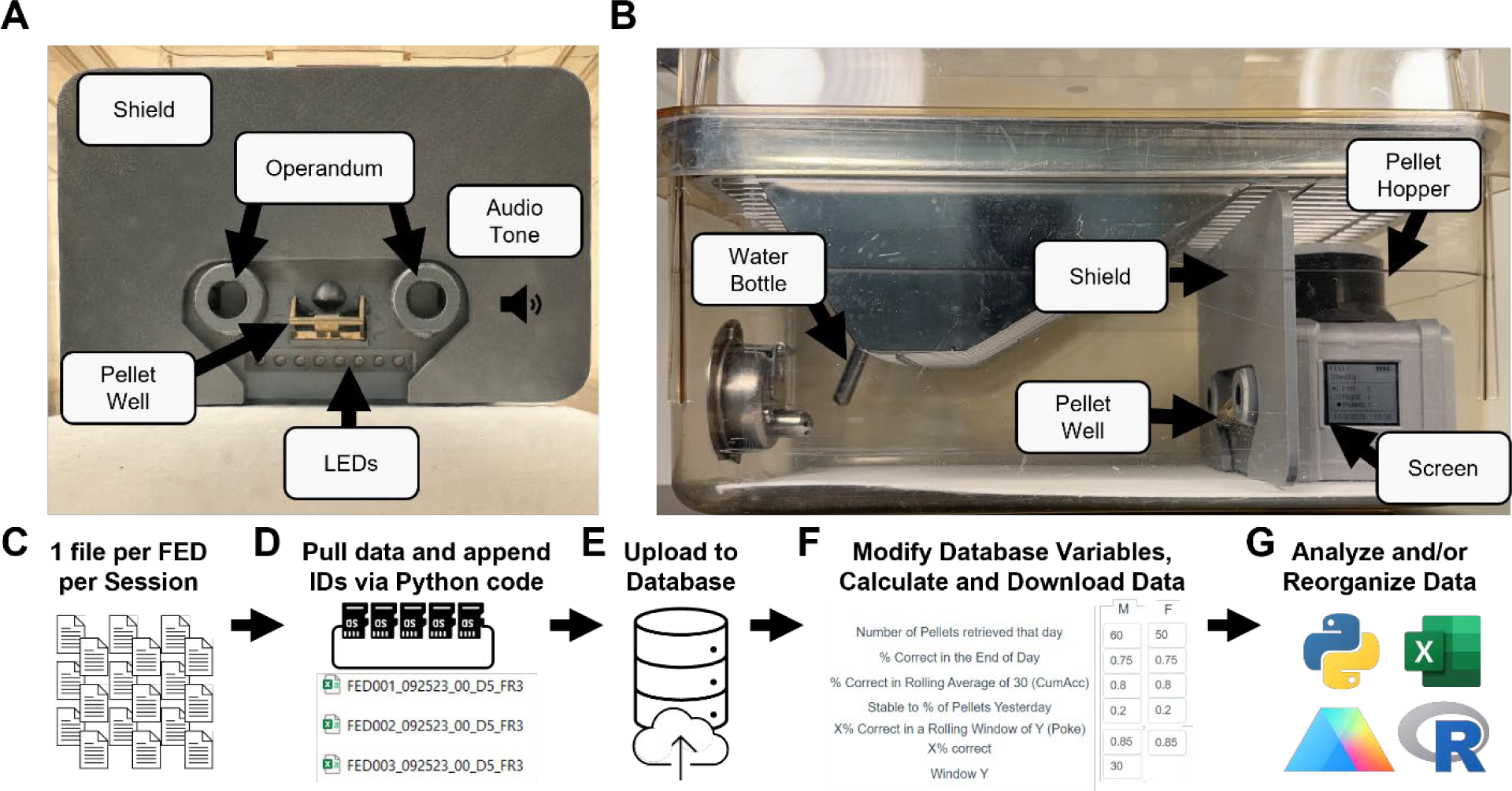
Description of FED3 setup and data analysis pipeline. **(A)** Front-view of the FED3 in a standard mouse home-cage demonstrating the main mouse-interaction features including two operandum, a pellet well, and LED and audio cues. **(B)** Side-view of FED3 in a standard home-cage demonstrating the additional features including the FED pellet hopper and screen, a custom-printed shield to prevent the mouse from climbing on the device, and the bottle for water consumption. **(C-G)** Data pipeline optimized for high throughput data cognitive behavioral analysis using the FED3 including: **(C,D)** bulk file collection from individual SD cards using Python code, **(E)** data upload to a Python-based database followed by **(F)** data analysis via user-defined experimental thresholds, and downloading of the compiled results as a CSV file for statistical analysis using preferred software. A detailed guide to the pipeline and all custom codes can be found in the Supplemental Information.

## RESULTS

The FED3 has been successfully implemented in non-operant quantitative feeding studies, but published operant studies largely focused on motivated consumption using fixed or progressive ratio testing procedures^52–55^. More complex cognitive behavioral testing often requires larger group sizes than traditional feeding studies to achieve significance, indicating a potential data management bottleneck for cognitive behavioral studies utilizing the FED3 where a 40-mouse study could generate over 1000 individual data files. Additionally, these studies show that mice perform fewer operant responses per hour (and thus receive fewer food pellet reinforcers) in a low-stress home-cage environment using the FED3 than they do using traditional operant behavioral apparatus^49,56^, indicating that utilizing the FED3 to evaluate cognitive behaviors will require optimization. Thus, we focused on addressing these two issues before initiating cognitive assessments using FED3.

### Data pipeline for cognitive behavioral assessments

To address data management when using the FED3, we developed an open-source, Python-based data pipeline optimized for performing cognitive behavioral testing (**Figure 1C-G**). First, the SD card from each FED must be removed. After connecting the SD cards to a computer using a multi-port USB hub, a custom Python-based code imports files indicated by a user-updated reference file. It then appends the imported file names with several identification codes which are used to indicate the subject, test day, and test type. Users then upload these appended files to a custom database where each subject is assigned a unique ID, allowing for detailed organization of files by subject by test day. Users can then input their preferred analysis and acquisition parameters from a range of researcher-defined options such as single or multi-day criteria, minimum number of pellets retrieved, minimum end-of-day percent correct, etc. The database then generates a CSV file containing the results of this analysis organized by subject and day, which users can download and reorganize for analysis via their preferred software. Using our semi-automated pipeline, we estimate that the complete data compilation and analysis for a 40-mouse study can be reduced to approximately 30 minutes per day. Furthermore, this approach can be scaled such that larger, high-throughput studies become feasible using FED3.

### Fixed ratio impacts multiple parameters of operant performance using FED3

To establish optimal FED3 parameters for assessing cognitive function, we investigated the effect of reinforcement schedules on body weight, food consumption, and operant performance (**Figure 2)**. Male C57BL/6 mice (8-10 weeks) were group housed and given group magazine training for two sessions using the free feeding program (FF), which allows continuous non-operant food access by dispensing a food pellet each time a pellet is retrieved. Before session 3, mice were divided into one of four groups. Group 1 (FF) continued to use the FF program for the remainder of the experiment, while Groups 2-4 began operant conditioning using a fixed ratio 1 schedule of reinforcement (FR1) for sessions 3-8. At session 9, group 2 remained in FR1 while Group 3 and 4 increased to FR3 and FR5, respectively, for the remainder of the experiment. For each daily session, mice were single-housed with their own FED3 for 8 hours before returning to the group-housed cage for 16 hours. Using these four groups of reinforcement, we evaluated multiple parameters that may impact behavioral outcomes (complete statistics results compiled in **Supplemental Table 2**).

**Figure 2.**
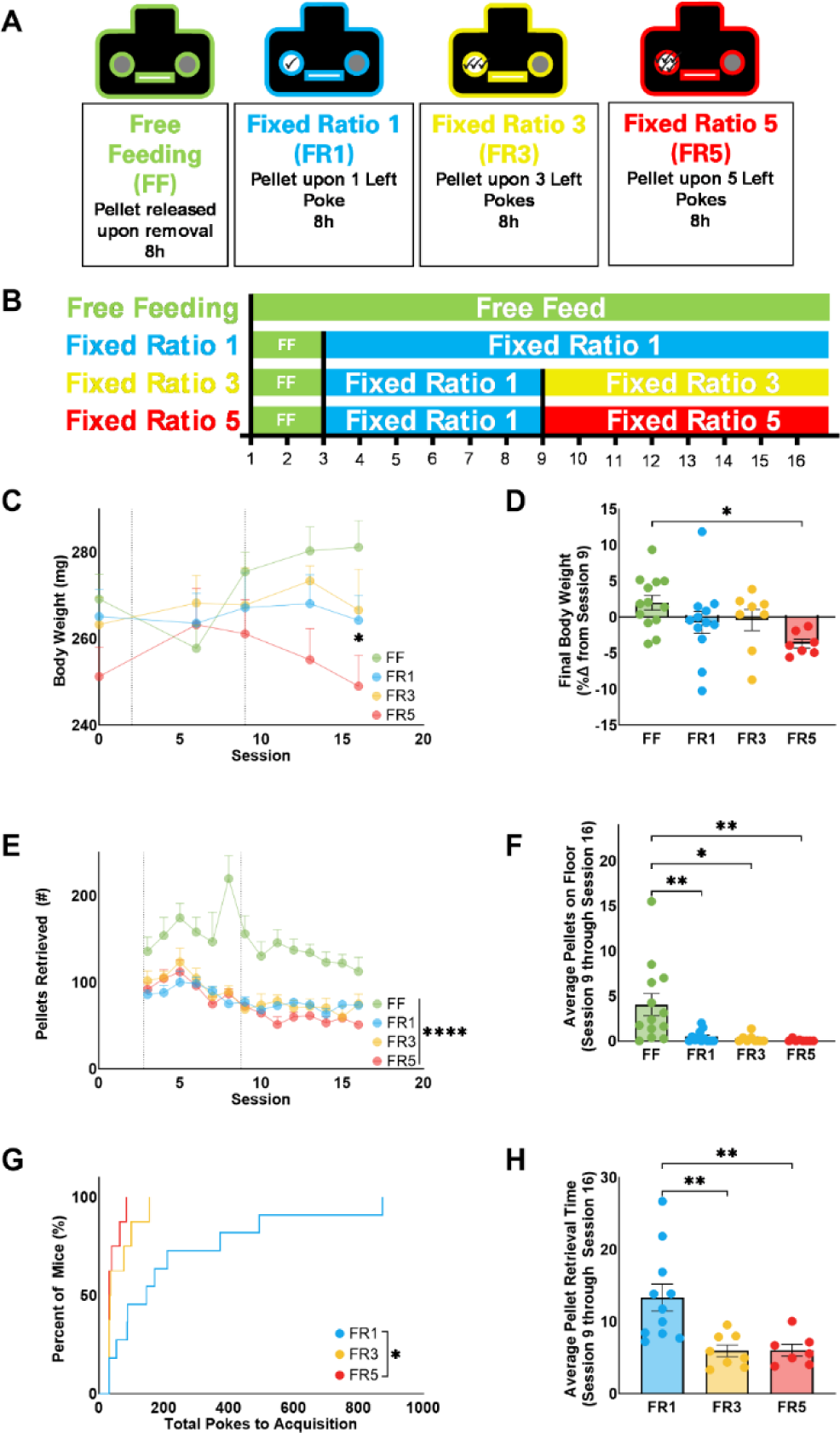
Fixed ratio impacts multiple parameters of operant performance using FED3. **(A)** Description of FED3 programs used to determine the optimal parameters for operant performance including Free Feeding (FF), Fixed-Ratio 1 (FR1), Fixed-Ratio 3 (FR3), and Fixed-Ratio 5 (FR5). **(B)** Experimental timeline for evaluating parameters of operant performance and goal-directed behaviors of four groups of male mice: FF (green, n=13), FR1 (blue, n=11), FR3 (yellow, n=8), FR5 (red, n=8). All four groups started with 2 sessions of group-housed FF before being single housed for FF, FR1, FR3, and/or FR5 as outlined for 14 additional sessions. **(C)** Mouse body weight (mg) was evaluated after session 7, 9, 14, and 16. **(D)** Bodyweight on session 16 displayed as the percent change from session 9, when the FR3 and FR5 groups transitioned to their final Fixed Ratio, respectively. **(E)** Number of Pellets dispensed by the FED3 per day. **(F)** Number of pellets found on the floor of the testing cage (average of sessions 9 through 16). **(G)** Kaplan-Meier curve of the total number of nose pokes required for animals to reach acquisition criterion: 85% correct in a rolling window of 30 pokes (starting at session 9). Average time to retrieve a pellet from the pellet well (average of sessions 9 through 16). Data displayed as mean ± SEM (except G). Statistical significance indicated by *P<0.05, ** P<0.01, ****P<0.0001.

The fixed-ratio schedule significantly altered body weight. While the FF group increased weight over the course of 16 sessions, FR5 mice had significantly lower body weight by session 16 (**Figure 2C**). Accordingly, mice undergoing FR5 for 8 sessions exhibited a decrease in relative body mass (approximately 5%) compared with FF (**Figure 2D**). These weight changes were not observed among the FR1 or FR3 reinforcement schedules. Moreover, these changes in body mass could not be fully explained by changes in overall food consumption, as the FR5 mice retrieved a similar number of pellets as FR1 and FR3 (**Figure 2E**) and left a similar number of pellets on the floor of the cage (**Figure 2F**). Thus, we determined that FR5 was not appropriate for long-term cognitive testing using FED3 due to potentially confounding factors associated with the FR5-associated weight loss.

The fixed-ratio schedule also had an impact on motivation and learning parameters. Compared to FR1, mice in the FR3 and FR5 groups required substantially fewer nose pokes to acquire discrimination learning (DL), with all mice in these two groups reaching our threshold in less than 200 nose pokes and within two sessions (**Figure 2G**). Moreover, the time between when a pellet is dispensed by the FED3 and retrieved by the mouse is significantly reduced under both the FR3 and FR5 schedules of reinforcement (**Figure 2H**). Collectively these results indicate that FR1 did not produce sufficient goal-oriented behavior, while FR3 and FR5 both produce similar increases in operant performance. Thus, the cognitive behavioral testing in subsequent studies was performed using a FR3 schedule of reinforcement.

### Validation of cognitive flexibility following bilateral lesion of OFC

To demonstrate the validity of using the FED3 for cognitive testing, we developed a 6-test operant battery to evaluate multiple aspects of cognitive behavior in mice (**Figure 3A**). Following two sessions of magazine training under group housing conditions and two sessions of individual daily FR1 sessions, male C57BL/6 mice (10-11 weeks) were tested in daily FR3 sessions until meeting DL acquisition criteria defined as meeting or exceeding 85% accuracy in a rolling window of 30 pokes^57,58^. Following acquisition, mice were tested for cognitive flexibility (Reversal Learning, RL), response to reward devaluation (via Quinine adulteration, QU), response to increased instrumental effort (Progressive Ratio, PR), response to cue devaluation (Extinction, EX), and response to cue re-valuation (Reinstatement, RE) over 20 8-hour sessions (**Figure 3B**). Following each test, mice re-established stable FR3 responding before beginning the next test. Importantly, these commonly used tasks investigate distinct but related aspects of cognitive and goal-directed behavior. Complete statistical results for these tests are compiled in **Supplemental Tables 3, 5-8.**

**Figure 3.**
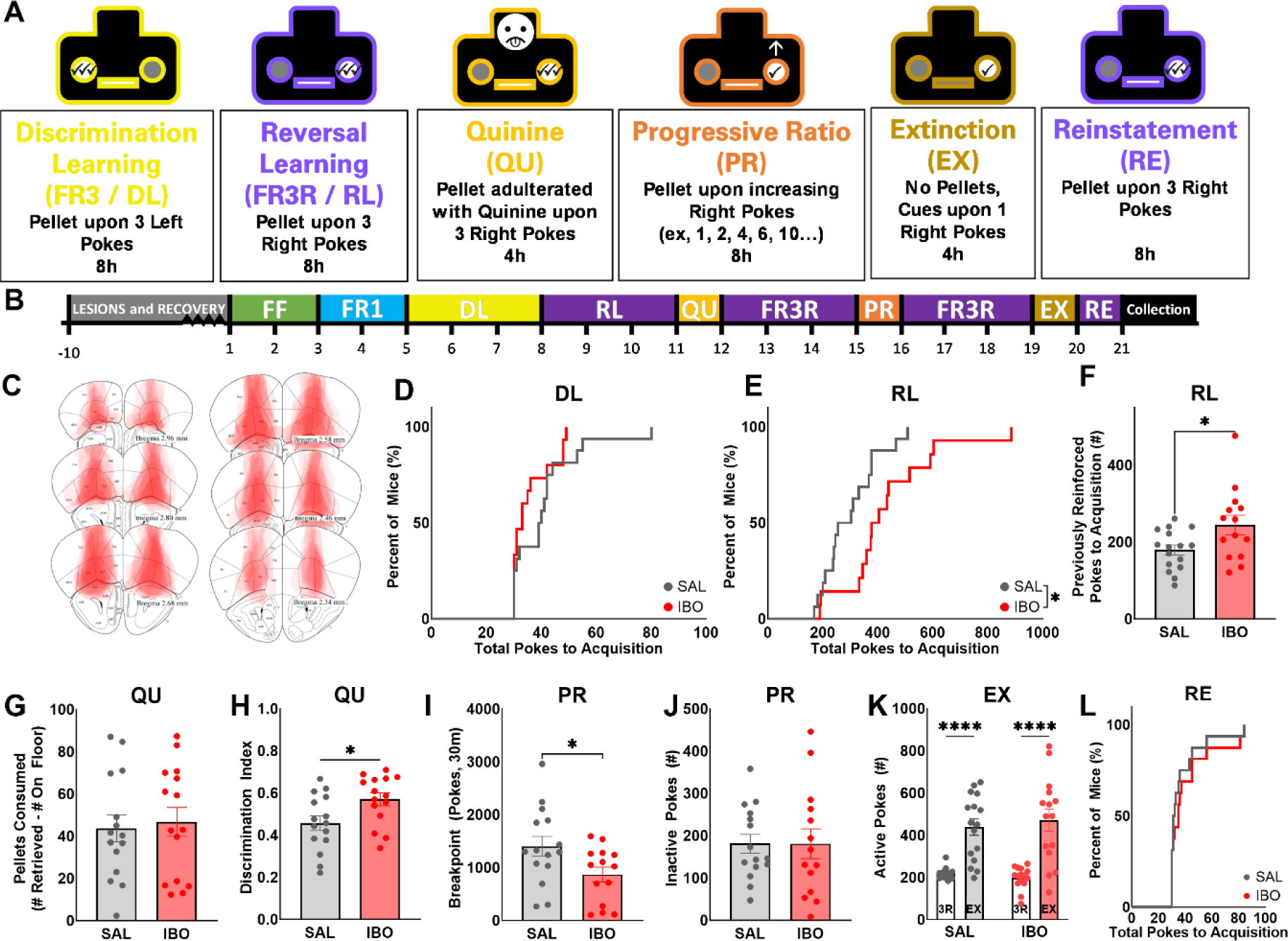
Bilateral OFC lesions reduce reversal learning behavior using FED3. **(A)** Description of FED3 programs used to evaluate cognitive performance including Discrimination Learning (DL, also FR3), Reversal Learning (RL, also FR3R), Quinine Test (QU), Progressive Ratio Test (PR), Extinction (EX), and Reinstatement (RE). **(B)** Experimental timeline for a typical mouse undergoing this behavioral battery, demonstrating lesion and recovery time followed by daily progression through tests. Male mice received bilateral injection of saline (SAL, n=16) or ibotenic acid (IBO, n=15) into the orbitalfrontal cortex (OFC), and were given 7 days to recover before initiating the 6 test behavioral battery. Tissue collection for histological verification of lesion efficacy and location occurred 27 to 32 days post-injection. **(C)** Schematic demonstrating lesioning of the lateral and ventral OFC across subjects, with each subject represented on a separate stacked layer. **(D, E)** Kaplan-Meier curve of the total nose pokes required for animals to reach the acquisition criterion of 85% during DL and RL. **(F)** Number of nose pokes on the previously reinforced nose poke until reaching acquisition criterion during RL. **(G)** Number of Quinine-adulterated pellets consumed during QU test. Consumption is calculated as pellets retrieved during the test minus pellets found on the floor. **(H)** Discrimination Index during QU test. **(I)** Number of active pokes made before reaching a breakpoint, defined as 30 minutes of inactivity during PR test, or final number of pokes. **(J)** Number of inactive pokes made during the PR test. **(K)** Number of active pokes made during prior four days of 3R (3R) or on the initial EX session demonstrating extinction burst. **(L)** Kaplan-Meier curve of the total nose pokes required for animals to reach acquisition criterion of 85% during Reinstatement (RE). Data displayed as mean ± SEM (except D, E and L). Statistical significance indicated by *P<0.05, ****P<0.0001. Diagrams modified from Paxinos and Franklin (2001).

To demonstrate that a subset of cognitive behaviors was driven by discrete cortical activity, we evaluated the effect of chemical lesioning the orbital frontal cortex (OFC) on our behavioral test battery. Before the experiment, mice received bilateral injections of saline (SAL) or ibotenic acid (IBO, an excitotoxic agent) into the OFC and recovered for 7 to 10 days **(Figure 3B)**. Immunohistological staining of the cortex performed post-study indicated that the lesioned regions consisted of large areas of the lateral and ventral OFC, but largely spared the medial and dorsolateral OFC regions with minimal impact on the medial prefrontal cortex (**Figure 3C**). While lesioned and sham mice needed a similar number of total pokes (**Figure 3D**) to acquire DL, the IBO-lesioned group required significantly more pokes than SAL controls to complete RL and made more presses on the previously reinforced nose poke (**Figure 3E-F**). Compared with DL, both groups required substantially more nose pokes to reach criterion during RL, suggesting that this test presents a challenge in the flexible adjustment of previously learned behavior (**Figure S1B**, **S1C**). Collectively, these results demonstrate that the FED3 captures the OFC lesion-induced RL impairments previously identified in both human and rodent models^59^ using traditional operant apparatus-based approaches.

The OFC is involved in numerous aspects of associative learning and goal-directed decision-making, including those related to reward valuation^60^. As such, we investigated the response to reward devaluation via the bitterant quinine in IBO-lesioned mice and SAL controls. Consistent with a role of the lateral and ventral OFC in cognitive flexibility and moderating the response to reward devaluation, but those regions not having a role in the perception or response to quinine, we find that IBO lesioned animals did not demonstrate altered quinine consumption (**Figure 3G**) but maintained a higher Discrimination Index (**Figure 3H**) during the test compared to SAL controls. Thus, while there were no significant differences between groups in the number of active or inactive pokes at the end of the test (**Figure S2C-D**) or by repeated measures analysis in 30-minute bins (**Figure S2F-G**), IBO lesioned animals demonstrated a significantly higher correct poke bias (**Figure S2H**) and end of day percent correct (**Figure S2E**), unlike SAL control animals who demonstrated non-significant increases in activity on the inactive noes poke in response to reward devaluation. Further consistent with the absence of a role of the OFC in innate taste aversion to quinine, both groups retrieved (**Figure S2A**) and left (**Figure S2B**) a similar number of pellets on the floor of the cage during the test. Collectively, the QU results suggest that lateral and ventral OFC lesions result in a failure to alter active poke activity in response to reward devaluation, absent changes in innate taste aversion, patterns of aversive reward retrieval, or aversive reward consumption.

Next, we investigated the effect of OFC inactivation on motivation and reward valuation. During the progressive ratio test (PR), IBO-lesioned mice made fewer active nose pokes (**Figure 3I**) and received fewer pellets (**Figure S3A**) than SAL controls before reaching breakpoint. However, there was no difference in the number of active or inactive pokes at the end of the day (**Figure 3J**) or to the end of day percent correct responding during the PR test (**Figure S3C**). Subsequent tests of reward valuation during extinction learning (EX) and reinstatement (RE) demonstrated that OFC-lesioned and SAL control mice exhibited similar active pokes, inactive pokes, and end of day responding on the active lever (**Figure S4A-C**) in a single-day EX test, which resulted in an extinction burst of increased activity compared to baseline performance (**Figure 3J**). Subsequent acquisition of operant performance for reward was similarly unaffected by treatment (**Figure 3K**). These findings indicate that the FED3 identified a role for the OFC in cognitive flexibility in response to increased instrumental effort.

### Implementation of cognitive testing in aged male and female mice

Many neurological disorders exhibit sexual dimorphism, with distinct underlying mechanisms and presentation of symptoms in males and females^61–66^. Furthermore, the symptoms for many diseases emerge later in life or are worsened by aging, and thus testing mice in early adulthood – when cognitive performance is most robust – may not be relevant to the disease model. To demonstrate that cognitive testing using the FED3 can address these concerns, we implemented our test battery to evaluate cognitive function in 9-11 month old mice of both sexes from a genetically modified mouse line predicted to impact cognitive behavior.

The FAAH P129T mutation (**Figure 4A**) has been identified in clinical populations as a predictive biomarker for problematic drug use^67,68^, and is implicated in pain tolerance^69,70^ as well as anxiolytic phenotypes^71^. Both humans and mice with this polymorphism exhibit dysregulation of corticoamygdalar circuitry^71^, however, the role in cognitive behaviors remains unexplored. Thus, we performed a complete cognitive assessment (**Figure 3B**) using 9-11 month old wild-type (WT) and FAAH P129T knock-in (KI) mice of both sexes. Using these four groups, we evaluated multiple goal-directed parameters that may impact behavioral outcomes (complete statistical results compiled in **Supplemental Tables 4, 9-12**).

**Figure 4.**
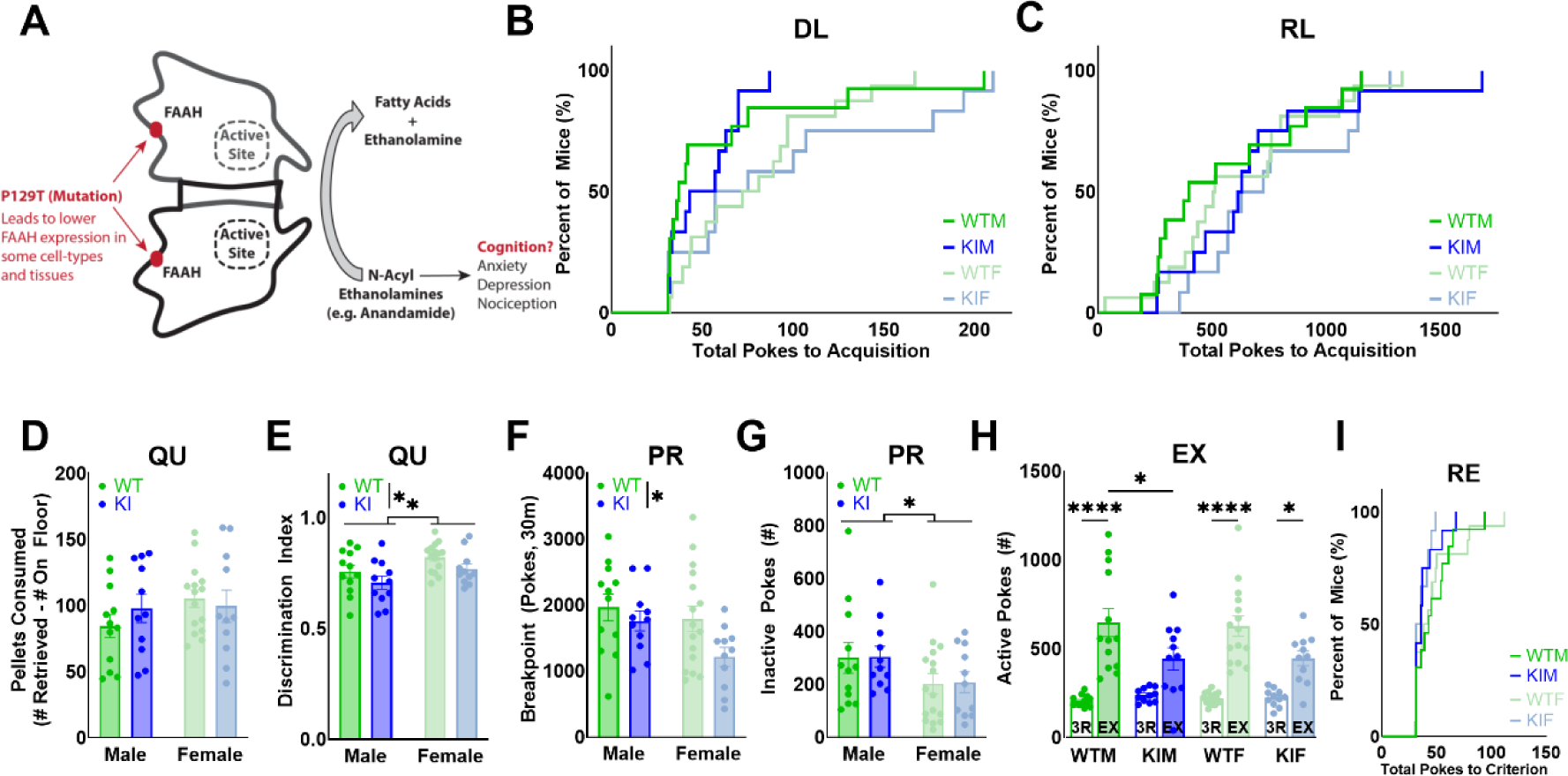
Both FAAH P129T genotype and sex influence cognitive performance in aged mice. **(A)** Fatty acid amide hydrolase (FAAH) plays a critical role in corticolimbic signaling and influence multiple behaviors including nociception, anxiety, depression, and cognition. The P129T single nucleotide polymorphism occurs outside the FAAH active site, yet it can dysregulate corticoamygdalar circuitry and increase problematic drug use in clinical populations. For this study, we evaluated wild-type (WT) and FAAH P129T knock in (KI) mice of both sexes for cognitive impairments using FED3. **(B)** Kaplan-Meier curve of the total nose pokes required for animals to reach the acquisition criterion of 85% correct in a rolling window of 30 pokes during DL. **(C)** Kaplan-Meier curve of the total nose pokes required for animals to reach the acquisition criterion of 85% correct in a rolling window of 30 pokes during RL. **(D)** Number of Quinine-adulterated pellets consumed during the Quinine (QU) test. Consumption is calculated as pellets retrieved during the test minus pellets found on the floor. **(E)** Discrimination Index during the QU test. **(F)** Number of pokes made before reaching a breakpoint, defined as 30 minutes of inactivity during the Progressive Ratio (PR) test. **(G)** Number of inactive pokes made during the PR test. **(H)** Number of active pokes made during prior four days of 3R (3R) or on the first Extinction (EX) session demonstrating extinction burst. **(I)** Kaplan-Meier curve of the total nose pokes required for animals to reach the acquisition criterion of 85% correct in a rolling window of 30 pokes during Reinstatement (RE). Data displayed as mean ± SEM where applicable from four groups: WT males (n=12-13), KI males (n=11-12), WT females (n=15-16), and KI females (n=11-12). Statistical significance indicated by *P<0.05, ** P<0.01, ****P<0.0001.

Our study identified distinct genotypic effects on cognitive behavior in mice. No genotypic effects were revealed during acquisition of DL (**Figure 4B, S5A**) or RL (**Figure 4C, S5B-F**). During the QU test we observed significant effects of genotype on Discrimination Index (**Figure 4E**) and end of day percent correct (**Figure S6E**) without changes in overall active or inactive (**Figure S6C-D**) pokes, suggesting that P129T KI animals may demonstrate increased behavioral flexibility in response to reward devaluation. There were no sex or genotype effects on quinine pellets eaten (**Figure 4D**), retrieved, or left on the floor (**Figure S6A-B**) suggesting these effects are not due to changes in the innate aversiveness of quinine. In response to increasing instrumental effort during the PR test, there was a main effect of genotype on the breakpoint as measured by active nose pokes (**Figure 4F**) and pellets (**Figure S7A**), with KI animals making fewer total active pokes than their WT counterparts **(Figure S7B)**, but no genotypic effect on inactive nose pokes (**Figure 4G**) or end of day percent correct (**Figure S7C**). On the first day of extinction, all groups increased active nose poke responding (extinction burst) compared with prior baseline FR3 performance once the FED3 stopped delivering food pellets, but the effect was stronger in WT animals **(Figure 4H).** As such, during the extinction burst, there was a main effect of genotype on the number of active **(Figure S8A)** and inactive (**Figure S8B**) nose pokes, with KI making fewer responses than WT mice. We further measured how long it took animals to extinguish learned behavior, however, there was no genotypic effect in the differences in the total number of nose pokes or sessions to reach extinction criteria **(Figure S8C-D)** or reinstatement (**Figure 4I**). We hypothesized that behavioral changes during PR and EX (but not DL/RL) could indicate dysregulation of other corticolimbic regions, such as the medial prefrontal cortex^72,73^ and the dorsal striatum, and using activity-based protein profiling we confirmed that P129T KI mice exhibited diminished FAAH activity at both of these brain sites, in addition to other brain regions broadly associated with reward, learning, and memory including the hippocampus, OFC, amygdala and VTA (**Figure S10**).

We also observed findings consistent with prior research indicating that aged animals demonstrate cognitive inflexibility as compared to younger animals by comparing performance in 9–11-month-old P129T WT males to the 10-11 week old Saline-injected C57BL6/J males. While age did not impact the total number of pokes required to reach acquisition of DL (**Figure S5G**), the aged P129T WT males required significantly more pokes to reach acquisition of RL (**Figure S5H**). Collectively, these findings demonstrate that the FED3 captures expected age-related cognitive impairmentsanddoes not preclude aged animals from completing the test battery in a timely manner.

Additionally, we also identified multiple sex-specific effects that suggest females may use different cognitive strategies during testing. No sex-specific effects were revealed during acquisition of DL (**Figure 4B**), RL (**Figure 4C**), nor did they impact the number of pellets consumed during the QU test (**Figure 4D**) or the breakpoint in the PR test (**Figure 4F**).

However, the females made more inactive nose pokes during DL (**Figure S5A**), and fewer inactive nose poke during QU (**Figure S5E**) and PR (**Figure 4G**) tests. While no effect of sex was measured on active (**Figure S8A**) or inactive (**Figure S8B**) nose pokes during the extinction burst, post hoc analysis revealed that the genotypic reduction of inactive nose pokes was driven largely by female KI mice. Collectively, these findings demonstrate that the FED3 can effectively compare cognitive tests in male and female mice using active nose pokes as an outcome measure, but sex-differences in inactive nose pokes during multiple tests may suggest that females may be less likely to alter habitual performance during some operant tasks.

## DISCUSSION

Evaluation of cognitive behaviors using operant tests remains a critical component for understanding and treating a wide range of neurobiological disorders^74–76^. By addressing technical challenges related to experimental parameters and data management, we demonstrated that the open-source FED3 can be scaled to run large cohorts of mice simultaneously in low-stress, home-cage-like environments. Using this approach, we demonstrated that a 6-test operant battery can be completed in as few as 20 sessions and measure multiple dimensions of cognitive function including associative learning, cognitive flexibility, reward valuation and devaluation, and motivation. We validated this approach using site specific inactivation of the OFC and used the FED3 to identify genotypic, age-, and sex-specific changes in cognitive behaviors.

Our study supports an important role for stress in the underlying mechanisms of cognitive behaviors^15,17^. In this study, mice have unrestricted access to receive their daily food requirements during extended daily sessions and show no weight loss during FR1 and FR3 reinforcement schedules when compared with the free feeding group. However, mice exhibited a loss of weight during FR5 reinforcement that could not be explained by food consumption alone, as these animals ate the same number of food pellets as FR1 and FR3 mice. This suggests that the increased work needed for mice to receive sufficient food pellets during longer sessions may result in changes to metabolic or neuroendocrine systems. Accordingly, mice treated with the stress hormone corticosterone exhibit weight loss despite consuming more food^77^. By comparison, animals undergoing food reinforcement during short time-constrained paradigms may also be influenced by stress and anxiety-like reponses^78–81^. Studies show that mice may request more rewards than they actually consume during short reinforcement sessions under food-restricted conditions^82,83^, suggesting that stress caused by perceived food instability may lead to observed hoarding behaviors and influence operant responding^84^. This stress may impinge on cognitive function as multiple lines of evidence support an inverted-U-shaped curve ^85^ for the influence of stress on learning and cognition, where moderate levels of stress enhance learning, while low or high stress impairs it. For instance, acute mild stressors increase performance during a paired associative learning task^86^, and increase performance during a RL test^87,88^. However, other studies have found that acute or chronic stress results in strategy shifts associated with a switch to habitual activity at the expense of goal-directed behavior^85,89,90^. In our protocol each animal has a dedicated testing cage they inhabit for 8h, which may encourage familiarity with the system and reduce the stress of testing. Thus, it is possible that a testing approach with reduced environmental stress may facilitate learning and executive function or be otherwise conducive to operant testing.

Our study using the FED3 demonstrates that the lateral and ventral OFC influences distinct aspects of executive functions. We found that inactivation of these regions of the OFC spared discrimination learning but impaired reversal learning performance^57,59,91^ as measured by total pokes and errors to acquisition, reflecting the well-documented role of the OFC in flexible decision-making and perseverative behavior during reversal. Additionally, lesioned animals demonstrated a lower progressive ratio breakpoint without impacting error rate (captured by inactive nose pokes) during this test, supporting a previously described effect of lateral OFC inactivation in reducing motivated effortful behavior^73^. As others have reported, OFC lesioning had no effect on the number of quinine-infused pellets retrieved or consumed indicating that innate taste aversions are not controlled by these regions^92^. However, lesioned animals demonstrated a resistance to altering activity during the test, maintaining an overall active poke bias and higher discrimination index. These results are in line with other studies which find that OFC lesioned animals continue to approach cues associated with devalued rewards^93^ and demonstrate a negative devaluation index^94^. Collectively, these results support the construct validity of our FED3 testing approach, demonstrating that it can be used to capture OFC-dependent behavioral changes. However, the OFC is also implicated in other aspects of decision-making broadly relating to value and outcome expectancies ^95–97^ and thus some studies report that inactivation of the OFC impairs extinction learning that were not seen in our study.

This difference may be explained by the amount of time between lesion and testing ^93,98^, as neuronal plasticity and other compensatory mechanisms can lead to functional recovery in mice after a few weeks that mask the impact of the OFC during behavioral testing. Consistent with this, we performed extinction testing at least 20 days post-lesion and it is possible that OFC inactivation at an earlier timepoint may produce different results. It is also possible that the inclusion of cues after an active poke during this test may make it harder to compare to extinction tests where a cue is presented before action to illustrate operandum activity. Future studies could be done to examine the effect of OFC lesioning on cue-driven behavior by investigating un-cued extinction, cue reinstatement absent reward, and finally cue and reward reinstatement, as prior studies have demonstrated that the OFC is implicated in cue-induced reinstatement of cocaine seeking^99,100^. Despite these limitations, we demonstrate that the FED3 can be used to capture aspects of executive dysfunction commonly associated with OFC inactivation.

Many traditional appetitive operant tests find sex-differences ^101–104^ in reward consumption or operandum activity, which can preclude their comparison. We found no significant sex differences in the total number of pokes to discrimination or reversal learning, indicating that this protocol may be useful to investigate the interaction of genotype and sex on executive function in mouse lines, as we did for the FAAH P129T mice. While we did not observe differences in cognitive flexibility, during this investigation we observed that males increased activity on the inactive nose poke in response to both reward devaluation and increasing instrumental effort, an effect not observed in females. These differences may indicate that males employ a different response strategy than females in response to reward devaluation and increasing effort. Indeed, prior research has found that female rodents form habitual behavior more quickly than males^102^, which may contribute to our observation that females maintained activity on the active poke in response to these tests. It is possible that stronger habitual activity or habit formation in females led to maintained performance on the active lever despite devaluation and increased effort, while males instead investigated whether the inactive lever may lead to typical reward dispensation.

In conclusion, we demonstrate that the FED3 can function as an effective in-cage tool for evaluating cognition and learning behaviors in mice. The low cost of this approach allows for high-throughput behavioral evaluations on a rapid timeline while maintaining the sensitivity of traditional approaches. To encourage broader implementation of this approach, we provided detailed methods for using a data pipeline as well as troubleshooting FED3 technical errors (Supplemental Table 1). Future studies will expand the variety and complexity of the behavioral tasks and apply these approaches to study the role of genetic and inducible models of neurological disorders that impact cognitive function.

## METHODS

### Reagents and Consumables

Materials were purchased as follows: **Behavioral Testing:** Braintree Scientific: Iso Pads, 6”x10” (#ISO); Bio-Serv: Dustless Precision Pellets, 20 mg, Rodent Purified Diet (#F0071); Dustless Precision Pellets, 20 mg, Rodent Purified Diet, Quinine (0.44% by weight) (#F07619). **Surgery:** MWI: Isoflurane (#NDC 13985-528-60): AbCam: Ibotenic Acid (ab146670-1001): Corning: Phosphate Buffered Saline, 1X (#21-040-CM). **Histology:** Millipore-Sigma: Xylenes (#534056); Cresyl Violet Acetate (#C5042); Eukitt® Quick-hardening mounting medium (#03989); Anhydrous Sodium Acetate (#S2889); Sigma Aldrich: Ultrapure Sucrose (#RES0928S-A102X), Sodium Hydroxide (#415413), Ethanol, Reagent Grade (#362808); Fisher Scientific: Acetic Acid, Glacial (#S70048); Corning: Phosphate Buffered Saline, 1X (#21-040-CM).

### Animals

For experiments using only wild-type mice, C57BL/6J male mice (n=83) were purchased from Jackson Labs at 8-11 weeks and acclimated for 5-15 days before starting experimental procedures. All mice of both sexes (n=53) for experiments studying the FAAH P129T mutation were bred in-house using heterozygous × heterozygous breeding pairs of *Faah P129T* knock-in mice kindly provided by Dr. Benjamin Cravatt^71^ (Scripps Research) and acclimated until the start of experimental procedures at 21 to 26 weeks of age. All mice were group-housed 2 to 5 per cage on a 12-hour reverse light cycle (21:00 on/09:00 off). Animals were given ad-libitum access to water, but food access was only given during daily 8-hour FED3 sessions or supplemental feeding (on weekends when testing was not performed) or in supplemental feeding as published^56^. Body weight was monitored twice weekly to ensure mice did not lose more than 15% of baseline body weight. All protocols and experiments were approved by the Virginia Tech (Blacksburg, VA, USA) Institutional Animal Care and Use Committee (IACUC).

### Feeding Experimentation Device 3 (FED3) Setup and Data Pipeline

#### FED3 Experimental Setup

Feeding and operant performance was measured using the open-source Feeding Experimentation Device 3 (FED3) ^49,50^. The FED3 is a battery-powered operant device that consists of two nose pokes for operant training, a pellet dispenser, LEDs for visual reinforcement, a buzzer for auditory reinforcement, and a screen for experimenter observation (**Figure 1A, B**). An internal microSD card logs interactions with the nose pokes and pellet well in real time for further analysis. Additionally, each device is outfitted with a 3D-printed shield to prevent animals from climbing on or behind the FED3 to prevent moisture from entering the device (**Supplemental Information**). Mice were tested in one 8h session per day, unless otherwise described in specific behavioral tests. During each test session, mice were single housed under red light in a standard cage containing a FED3, an empty food rack holding a water bottle, and an IsoPad for bedding (traditional bedding can enter the pellet well and interfere with data collection). At the end of each test session, mice were group housed in a standard home-cage with traditional bedding.

#### FED3 Data Pipeline

Data collection and analysis for cognitive testing was streamlined using a Python-based data pipeline (**Figure 1C**). Session data was simultaneously retrieved from multiple SD cards using a multi-port USB hub equipped with micro-SD card readers using Python code (**Supplemental Information**) that utilized a reference file to append each FED3 data file name with mouse and study identifiers. Next, these files are uploaded to our custom Python-coded FED3 cognitive analysis database that allows users to modify acquisition criteria parameters as needed. Acquisition criterion can be set as meeting multiple criteria per day (number of pellets retrieved, stability to the previous day’s number of pellets, end-of-day percent correct activity, etc.) or as reaching a minimum percent correct in a rolling window of pokes. The database will organize and calculate the data based on these user-set parameters, the results of which can be downloaded as a CSV file that can be further analyzed using traditional data analysis software (e.g. GraphPad Prism). All code described in the FED3 Data pipeline is freely available via GitHub https://github.com/mwblab/fed_database for implementation and further customization.

### Cognitive Behavioral Procedures and Tests

The FED3 delivers pellet rewards based on user-defined programs written in Arduino. In the programs used for this study, all procedures and tests from a single FED3 record and time stamp the following parameters in a CSV file: active nose pokes, inactive nose pokes, pellets delivered, pellets retrieved, pellet retrieval time. The number of pellets found on the floor of each cage were determined by experimenter observation and included in the final data analysis.

#### Free Feeding

Under this program, the FED3 delivers a pellet immediately after a pellet is removed from the pellet well. This results in continuous access to a pellet independent of operant responding and can be used for magazine training to acclimate mice to the FED3 and the food pellets^49^.

#### Fixed Ratio Testing (FR)

For all FR testing, every n^th^ correct response (non-chaining) was paired with a pellet delivery, audio tone (1 sec), and a visual LED cue (1 sec). The left nose poke was coded as the active lever for FR1, FR3, and FR5 testing; the right nose poke was coded as the active lever for FR3 reverse (FR3R). Acquisition of FR behavior was defined as reaching or exceeding 85% accuracy in a rolling window of 30 pokes (e.g., 26/30 pokes correct). These criteria were based on other long-access, in-cage operant feeding experiments^57,58^. The percent of correct pokes, the time until acquisition, the number of pokes (and pellets) until acquisition were calculated. While we did not investigate poke bias in these experiments, in other experiments where we collect poke data during extended sessions of FF we observe that daily poke bias is normally distributed, with the majority of sessions ending with 0.4 to 0.6% preference for the right poke (**Figure S9A**). We further find that during FF side preference can vary greatly by day, likely indicating that animals do not display strong preference for either poke (**Figure S9B**).

#### Discrimination Learning (DL) and Reversal Learning (RL)

DL was performed using a FR3 reinforcement schedule (active nose poke = left), and acquisition of DL behavior was defined as exceeding 85% accuracy in a rolling window of 30 pokes (e.g., 26/30 pokes correct). After meeting criterion, mice maintained the FR3 schedule for at least 2 additional sessions before moving on to the next test. The percent of correct pokes, the time until acquisition, the number of pokes (and pellets) until acquisition were calculated. RL was performed identical to DL, except that a FR3R reinforcement schedule (active nose poke = right) was implemented.

#### Quinine Test (QU)

QU test was performed using a FR3R reinforcement schedule with grain pellets containing quinine. This test was performed in a single 4h session, followed by access to standard chow for 4h. In the following session, mice returned to 3R and re-reached acquisition criterion before starting the next behavioral test. In addition to all previously described parameters, the number of pellets consumed was calculated as the number of pellets retrieved minus the number of pellets on the floor of the cage at the completion of the QU test. Discrimination Index was calculated as (active nose pokes – inactive nose pokes)/(active nose pokes + inactive nose pokes)^105^.

#### Progressive Ratio Test (PR)

PR was performed using an escalating reinforcement schedule for grain pellets over a single 8-hour session. The number of responses on the active lever needed for reward dispensation increased exponentially based on the following equation: (ratio = ratio + round ((5 * exp (0.2 * PelletCount)) - 5))^49,106^. Breakpoint was defined as the highest number of reinforcers earned before a 30-minute break between pokes, or at end of session if no break of at least 30-minutes occurred^107^. In the following session, mice returned to F3R and re-reached acquisition criterion before starting the next behavioral test. In addition to all previously described parameters, the breakpoint and number of active/inactive nose pokes at the breakpoint were calculated.

#### Extinction (EX)

EX was performed using a FR1 reverse reinforcement schedule (active nose poke = right) in 4h sessions, with active nose pokes paired with audio and visual cues but not food pellet. Following each 4h EX session, mice were given 8-hours of access to standard chow before returning to the fasted group-housed home-cage for 12 hours. Extinction burst was defined as the behavior that occurs during the first EX session. Acquisition of EX behavior was defined as ending a session with active pokes ≤20% of the average number of active pokes from the pre-extinction baseline (average of the preceding four FR3R sessions).

#### Reinstatement (RE)

RE was performed using a FR3R reinforcement schedule (active nose poke = right), and acquisition of RE behavior was defined as exceeding 85% accuracy in a rolling window of 30 pokes (e.g., 26/30 pokes correct). The percent of correct pokes, the time until acquisition, the number of pokes (and pellets) until acquisition were calculated.

### Intracranial Bilateral Lesions

#### Surgical Procedures

Animals were anesthetized with isoflurane via a low-flow vaporizer (Somnosuite, Kent Scientific) induced at 5% and maintained at 1-3% concentration. Animals were then secured in a stereotaxic frame (David Kopf Instruments) via ear bar. The skull was exposed and a Dremel was used to create holes in the skull at the injection site. Injections were made with a 10-µl NanoFil Syringe (World Precision Instruments, Germany) with a 33 GA blunt needle (World Precision Instruments, Germany) mounted to a syringe pump (Pump 11 ELITE Nanomite, Harvard Apparatus) with an injection rate of 0.1 µl per minute with a 4-5 minute resting period before removal. Orbitofrontal cortex (OFC) lesions (AP: +2.55mm, ML: ±1.2mm, DV: +2.40mm from Brain OR +2.6mm from Skull) were made using established coordinates^108^ in 26 mice by bilaterally injecting 0.4uL of ibotenic acid dissolved in 80:20 PBS and 1M NaOH. 17 sham-lesioned mice underwent the same procedure but were injected with sterile PBS. Mice were given 7-10 days to recover before beginning the experiment.

#### Histology

Mice that received OFC lesions were evaluated for accuracy and precision of lesion placement. Briefly, mice were sacrificed via cervical dislocation and decapitation for brain collection. Their brains were removed and drop-fixed in 4% w/v paraformaldehyde in PBS overnight at 4°C, followed by up to 36 hours each in a gradient of 10%/20%/30% Sucrose in PBS. Brains were coronally sectioned at 50um on a cryostat, mounted on glass slides, and stained with cresyl violet. Briefly, sections were mounted to slides and allowed to dry overnight. The following day sections were dehydrated in Xylene (10 m), rehydrated in 100%, 95% and then 70% ethanol (EtOH) followed by deionized distilled water (3 m each). Sections were then stained in 1% cresyl violet solution (8 to 16 m), dehydrated in 70% and 95% EtOH (1 to 2 m each), followed by saturation in Xylene (30+ m) before mounting with mounting medium and cover glass. Lesion size and location were examined under microscope and with images captured by 5MP USB digital camera (MU500, AmScope). Mice showing off target injection tracks or showing less than 50% lesioning of the lateral and ventral OFC on one or both sides of the brain were excluded from the study.

#### Statistical Analysis

Statistical analyses were performed using GraphPad Prism (version 10.0.2). Data reported as mean ± SEM where applicable. Body weight over time was analyzed using two-way RM ANOVA followed by Tukey’s post hoc, with ANOVA statistics and significant post hoc p-values reported in **Supplemental Table 2**. Number of pellets retrieved by session was analyzed using Mixed-Effects Model (REML), as hardware failures resulted in occasional data loss, with no follow-up post hoc tests. ANOVA statistics are reported in **Supplemental Table 2**. Single variable tests between more than two groups (final body weight by reinforcement schedule, average pellets on floor by reinforcement schedule, etc.) were analyzed using one-way ANOVA followed by Bonferonni post hoc when appropriate with ANOVA statistics reported in **Supplemental Table 2**. Single variable tests between two groups (pellets consumed by treatment group, active pokes by treatment group, etc.) were analyzed using unpaired t-Test, with statistics and p-values in **Supplemental Table 3**. Single variable tests investigating sex × genotype effects (pellets consumed by sex × genotype, active pokes by sex × genotype, etc.) were analyzed using two-way ANOVA and Bonferroni post hoc when appropriate, with ANOVA statistics and relevant post hoc p-values reported in **Supplemental Table 4**. Kaplan-Meier learning curves (pokes to acquisition during DL or RL) were analyzed using Log-Rank (Mantel-Cox) for differences between all groups. If significant differences were observed for experiments with more than two groups, pair-wise comparisons were made between each. In such cases the p-values reported (**Supplemental Table 2**, **Supplemental Table 4**) have been multiplied by the number of comparisons made to account for Bonferroni’s correction. Device failures during a single day test (QU, PR) resulted in either re-testing the mice during a new session or removing the mice from the final analysis. Statistical outliers were determined using Grubb’s test across all related measures. In the event of multiple outliers within the same group, the individual with the higher z-value was removed.

## Abbreviations

Discrimination Learning (DL), Reversal Learning (RL), Discrimination and Reversal Learning (DL/RL), Ethanol (EtOH), Fatty Acid Amide Hydrolase (FAAH), Feeding Experiment Device 3 (FED3), Free Feeding (FF), Fixed Ratio (FR), Fixed Ratio 1 (FR1), Fixed Ratio 3 (FR3), Fixed Ratio 3 Reverse (FR3R), Fixed Ratio 5 (FR5), Heterozygous (HET), Wild-type (WT), Knock-in (KI), Ibotenic acid (IBO), Medial prefrontal cortex (mPFC), Orbitofrontal cortex (OFC), Phosphate buffered saline (PBS).

## Notes

### Competing Interest Statement

The authors have declared no competing interest.

